# XePhIR: The zebrafish Xenograft Phenotype Interactive Repository

**DOI:** 10.1101/2021.09.13.460043

**Authors:** A. Groenewoud, G. Forn-Cuní, F.B. Engel, B. E. Snaar-Jagalska

## Abstract

Zebrafish xenografts are an established model in cancer biology, with a steadily rising number of models and users. However, as of yet, there is no platform dedicated to standardizing protocols and sharing data regarding zebrafish xenograft phenotypes. Here, we present the Xenograft Phenotype Interactive Repository (XePhIR, www.xephir.org) as an independent data sharing platform to deposit, share and repurpose zebrafish xenograft data. Deposition of data and publication with XePhIR will be done after the acceptation of the original publication. This will enhance the reach of the original research article, enhance visibility, and does not interfere with publication or copyrights of the original article. With XePhIR, we strive to fulfill these objectives and reason that this resource will enhance reproducibility and showcase the appeal and applicability of the zebrafish xenograft model.

## Introduction and purpose

Since the transplantation of human metastatic melanoma cells into zebrafish blastula-stage embryos (*Danio rerio*) by Lee et al. in 2005^1^, the popularity of the zebrafish to model human cancer has been on a steady rise (Figure 1): scientists all over the world incorporate zebrafish as a model for cancer compound screening and utilize the zebrafish xenograft model to address basic questions in tumor biology and metastasis^2,3,4^.

**Figure 1.**
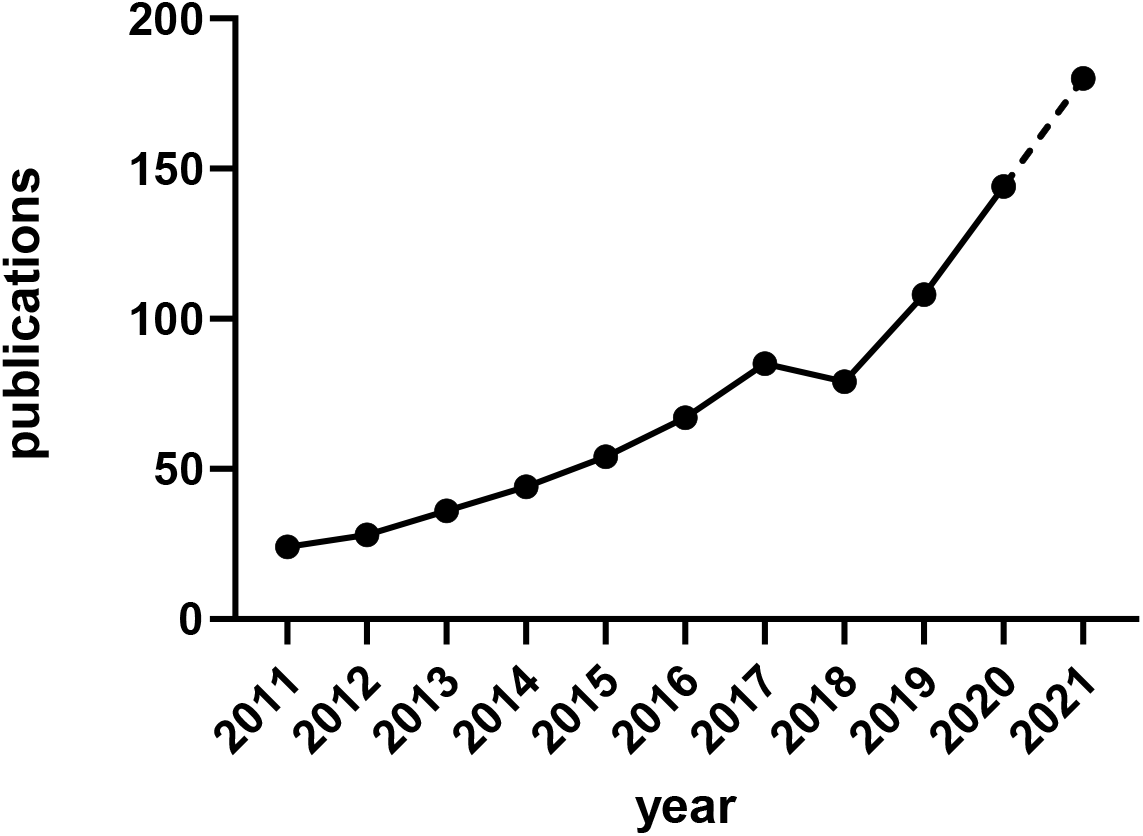
Increasing popularity of the zebrafish xenograft model. Over the course of the last 10 years the popularity of zebrafish xenograft models has been steadily increasing.

In 2016, we have published the first description of a zebrafish patient-derived xenograft^5^. One of the main issues we noted since then was a general lack of conformity among different laboratories working with zebrafish xenograft models (i.e., cell types, growth medium, cancer cell processing prior to engraftment, staining of cells, injection site and time). Recently, several highly translational studies using both stable cell lines and patient-derived xenografts have further supported the overall value of the zebrafish as a cancer model^6–8^. One of the main difficulties in evaluating and comparing data obtained with the zebrafish xenograft model as well as interlaboratory adoption is the lack of standardized protocols. Detailed protocols are needed due to the variable nature and needs of each cancer line, which may result in invalid engraftment phenotypes or incorrect data interpretation.

Considering the increasing interest and number of zebrafish-based studies in the cancer field and the ever-growing need for transparent and reproducible science, we have created a platform where zebrafish xenograft phenotypes can be showcased, in an open access database. This platform will help to exchange experience inside the zebrafish community as well increase visibility and appreciation for this useful cancer model by wider audience. The need for a platform of this kind is also evidenced by the recent initiative by Targen et al. 2020, who created a zebrafish xenograft metadata repository^9^. However, zebrafish research is often driven by image-based quantification analyses. Therefore, we believe that a visual data repository is a more fitting representation of the underlying data and furthers the goal of enhancing the reproducibility of zebrafish assays. We want to enable external users to quickly choose the cell lines and zebrafish models for their assay of choice, such as dissemination, orthotopic/ ectopic engraftment, angiogenesis, etc. (Figure 2). Alternatively, this platform will facilitate the comparison of phenotypes, cell lines and zebrafish models between labs to enhance the overall reproducibility of zebrafish models around the world. In addition, our approach enables easy outreach to users unfamiliar with the zebrafish xenograft model.

**Figure 2.**
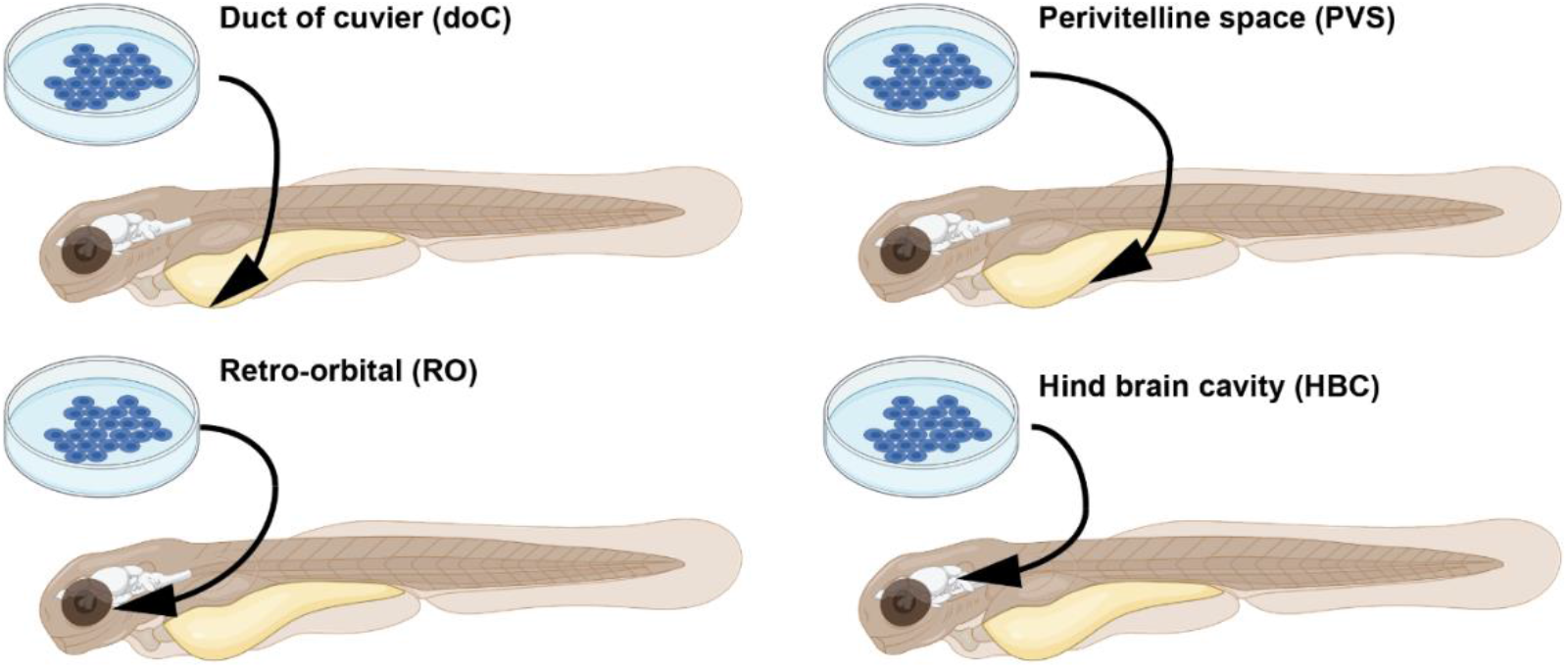
Graphical representation of common injection sites of the zebrafish xenograft model. Injection through the duct of Cuvier, a common route of injection to generate hematogenously disseminated cancer, also described as an experimental micro metastatic model.^10,11^ Perivitelline space injection, originally described as a model for the generation and assessment of angiogenesis, more recently developed as a model for the generation of primary-like tumors.^6^ Retro-orbital engraftment, used for the generation of orthotopic primary-like tumors derived from eye tumors, allows for the development of distant metastases.^12^ Hind brain cavity injection models, used for the generation of orthotopic brain cancer models and for the generation of brain metastasis models.^13^

Since most zebrafish assays are image-based, there is an untapped host of data, outlining the zebrafish phenotypes generated during the conventional course of the experiments. We therefore propose to use one representative image per timepoint for submission to XePhIR. Citations to the original research will be included to allow and stimulate the citation of the originator of the model to further enhance visibility.

### Comprehensive xenograft metadata and phenotype

To enhance the outreach potential of XePhiR, we will include a description of the cancer cells and explanation of the general experiment (a summary of the goal of the experiment and the outcome <150 words). To facilitate comparison and exchange between laboratories, we will include a comprehensive list of all metadata required to repeat the experiment (cell types, culture conditions, injection space, etc. as specified in Table T1) as well as phenotypes to check if well engrafted. Among the metadata, there are for example American Type Cell Culture (ATCC) and/or Expasy (Cellosaurus) identifiers for the engrafted cells, Addgene identifier for reporter constructs used, chemical tracer used (specified, by supplier and catalog number), and zebrafish line as by ZFIN identifier.

**Table T1:**
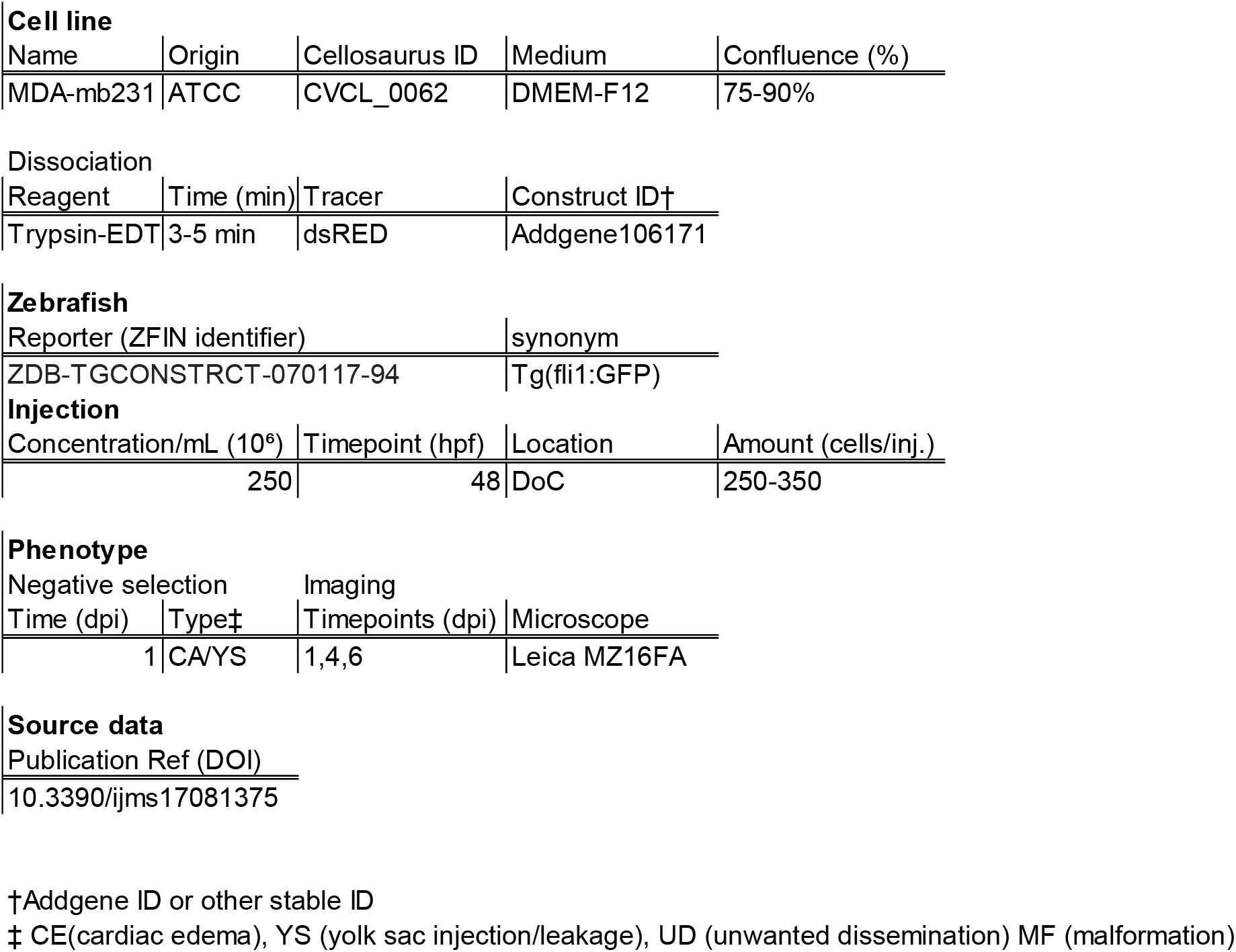
Example of the required metadata ensuring reproducibility and tractability of the published zebrafish xenograft models.

## Database contents

Since most zebrafish xenograft experiments are image-based, we have chosen for a graphic driven database. Here, users can upload one data set per timepoint of the performed zebrafish experiment (Fiji/bioformats compatible raw data only)^14^. The visual data will be supplemented with a comprehensive set of metadata as described previously (representation of the final website user interface in Figure 3).

**Figure 3.**
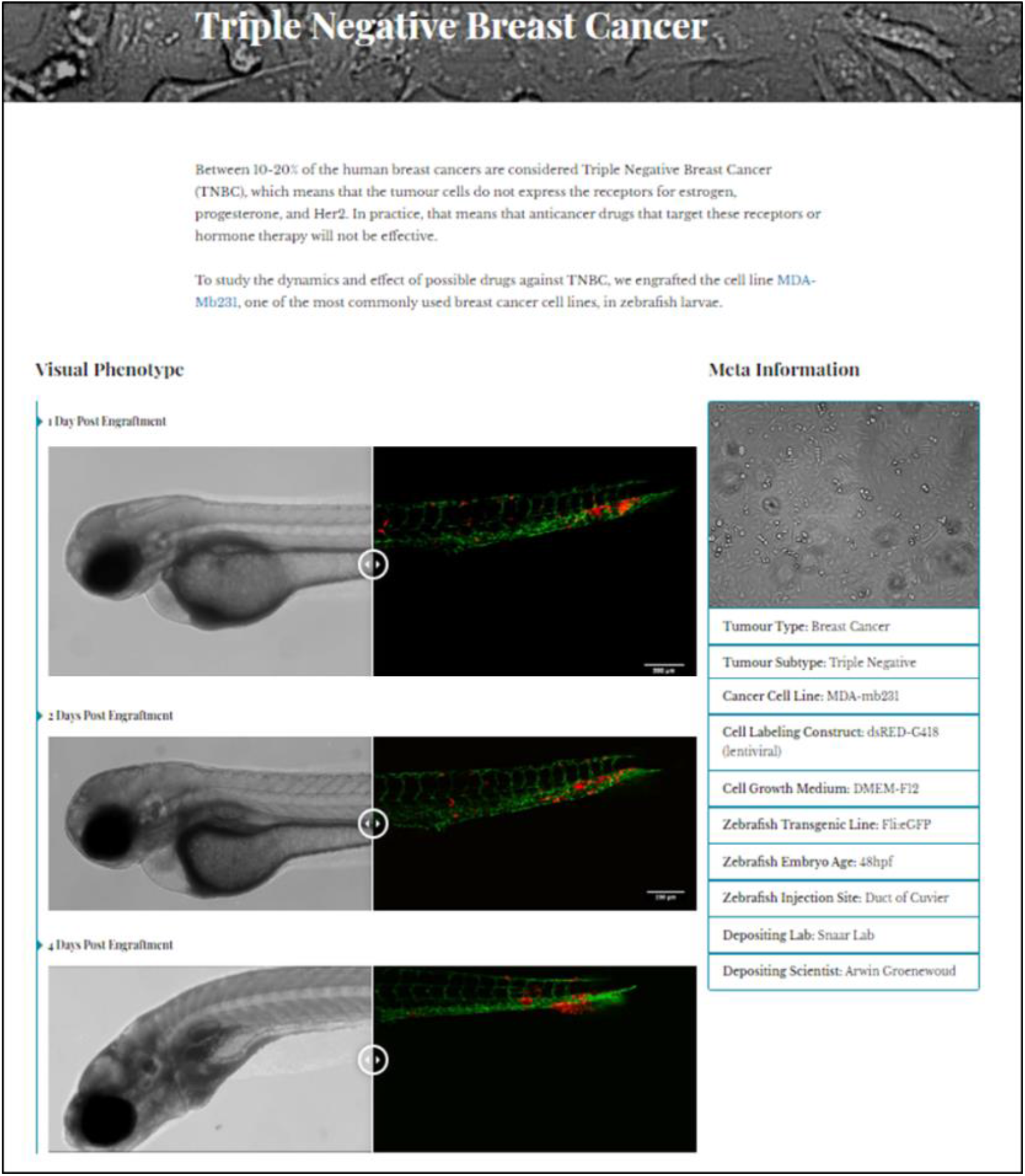
Triple negative breast cancer sample data as deposited. Image-based analysis data is used to generate a graph for the original publication and is subsequently uploaded to XePHIR (either directly or indirectly). Subsequently the available metadata is entered during submission (filling in the available data in the submission template). All metadata will be displayed next to the images that have been uploaded, when uploaded via Zenodo a digital object identifier will be provided to allow easy access to the original dataset.

### Standardized protocols

As to further enhance the ease of interpretation of protocols and to support the worldwide repeatability of zebrafish xenograft experimentation, we have provided standardized engraftment protocols. These protocols are dynamic and can be adapted through drop down menus to encompass all conventional injection sites (duct of Cuvier, perivitelline space, retro-orbital, and hind brain cavity), carrier solutions, cell densities, cell number, and time of injection (in hours post fertilization, hpf).

## Submission guidelines and Intellectual Property

Our database is open for submission. The focus of the submissions will be placed on the dissemination of already published data, to ensure that there will be no conflicts of interest between publishers and researchers. Next to the dissemination of previously published work, we will also provide a platform for unpublished data. Given that this could lead to a possible conflict of interest during later publication in a journal, this can only be done at the users informed request. If needed, uploaded data will be time-gated prior to publication, where the items will be released to the public after publication of the original manuscript.

To submit data to XePHIR, data has to be placed under a selection of creative commons licenses, namely the choice between the BY-NC-ND, BY-NC-SA and BY-SA licenses (more information at creativecommons.org). All CC licenses chosen disallow respective commercial use. Through CC licensing we can facilitate the use of the deposited material for future use in for instance grant applications. To do so, data can be uploaded either directly to XePHIR.org or via Zenodo^15^ as a whole dataset, thus automatically placing it under CC license and generating a digital object identifier (DOI) allowing the referencing of the data (Figure 4).

**Figure 4.**
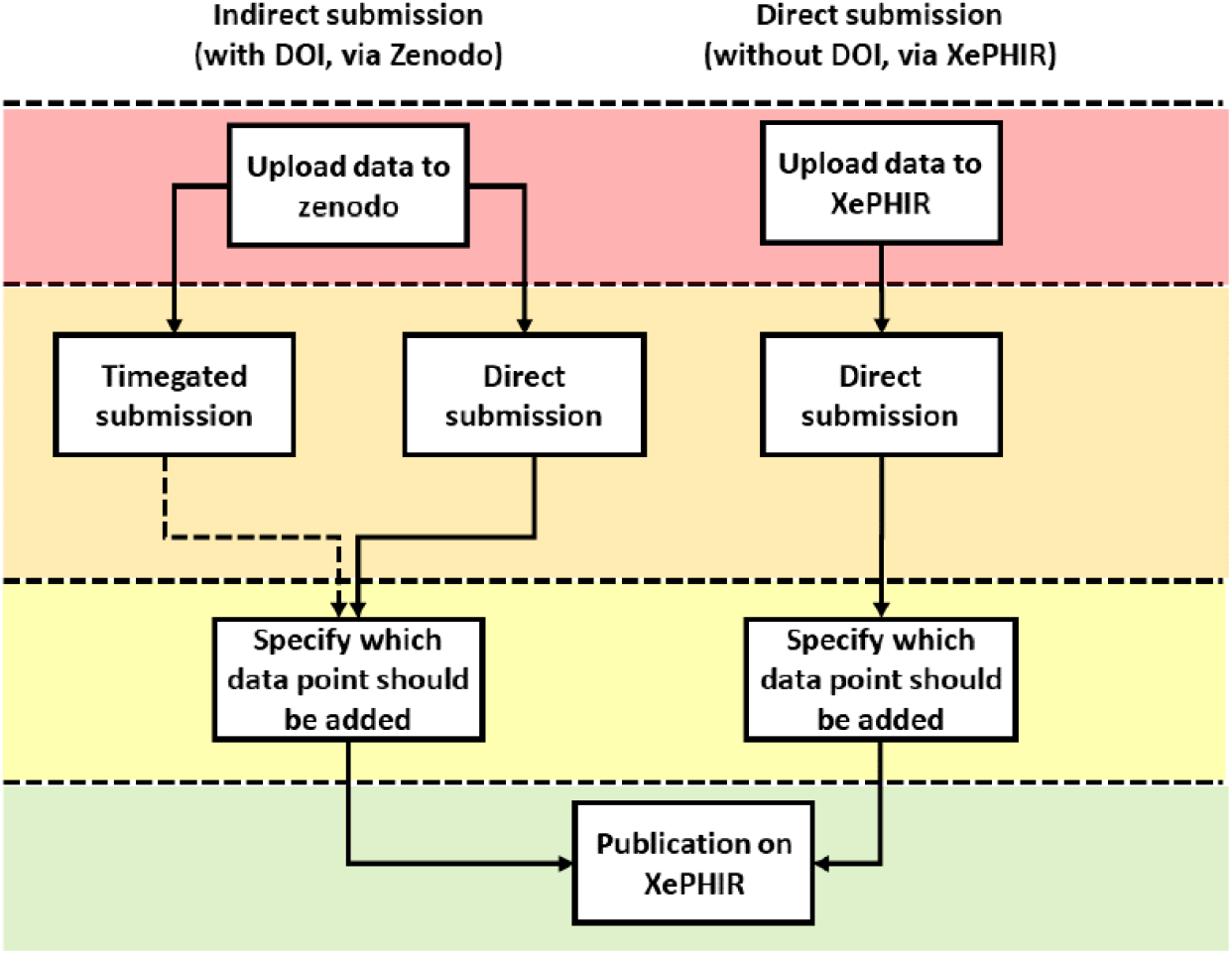
Methods of data submission to XePHIR. Indirect submission through Zenodo, allowing uploading of whole data sets (after acceptation of publication) will allow for enhanced transparency and re-usability of data and will provide the user with a DOI enabling citation of the data set. Uploading to Zenodo will automatically place the data under a creative commons license (CC). Direct submission to XePHIR will not provide the user with a DOI and will require the user to place the data under a CC license.

Careful selection of the representative data submitted to XePhIR will ensure a representative measure of the phenotype depicted and will thus enhance global reproducibility and minimize bias when selecting cancer cell lines for future experiments (Figure 5).

**Figure 5.**
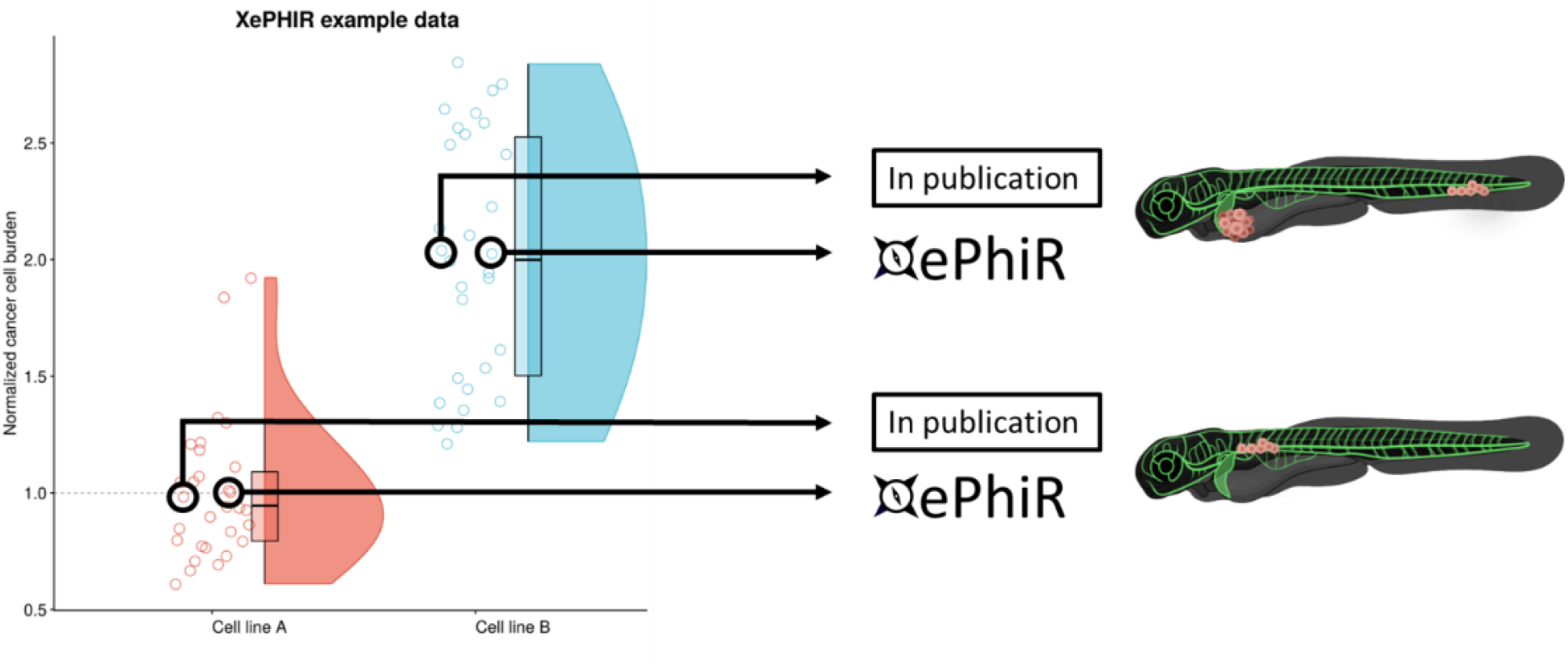
Data is derived from redundant image-based analysis. Image-based analysis data is used to generate a graph and is subsequently used for the publication of the original paper, where one sample individual is used in the original manuscript. Another representative individual is submitted to XePHIR either directly, or indirectly through deposition of the underlying dataset in Zenodo and subsequent publication of the representative image in XePhIR.

As the guidelines for the ethic use of animals vary between different countries, we will ask all depositors to sign a waiver stating that the generation of all data deposited has been generated in accordance with local animal ethical guidelines. We do not condone and take no responsibility for the unethical use of animals.

## Conclusion

With XePHIR we represent for the first time, a visual, user-driven zebrafish xenograft database which provides 1-page summaries with the most important information of a cancer study utilizing zebrafish (Figure 5), collating all required data to reproduce the experiments, referring to the original paper. Through XePHIR we will enhance the visibility of individual research groups and the zebrafish xenograft models they generate and use. Using a two-pronged data submission approach, we will be able to generate data for future comparative analysis between groups, models or cell lines, further enhancing the reproducibility of zebrafish xenograft models worldwide. Data submission through Zenodo would further enhance the data re-use capacity and allows for the referencing of individual (un)published data sets using the Zenodo generated DOI.

